# STIPS algorithm enables tracking labyrinthine patterns and reveals distinct rhythmic dynamics of actin microridges

**DOI:** 10.1101/2024.06.06.597299

**Authors:** Rajasekaran Bhavna, Mahendra Sonawane

## Abstract

Tracking and motion analyses of semi-flexible biopolymer networks from time-lapse microscopy images are important tools that enable quantitative measurements to unravel dynamical and mechanical properties of biopolymers in living tissues important for understanding their organization and function. Biopolymer networks pose tracking challenges as they exhibit continuous stochastic transitions in the form of merges/splits resulting in local neighborhood rearrangements over short time/length scales. We propose the STIPS algorithm (*S*patio*T*emporal *I*nformation on *P*ixel *S*ubsets) that tracks merging/splitting events in self-organizing patterning systems, by creating pixel subsets to link trajectories across consecutive frames. We demonstrate our method on actin-enriched protrusions, the ‘microridges’ that form dynamic labyrinthine patterns on outer surfaces of squamous cell epithelia, mimicking ‘active Turing-patterns’. We uncovered two distinct actomyosin based rhythmic dynamics within neighboring cells; common pulsatile mechanism between 2-5.9 mins period governing both fusion and fission contributing to pattern maintenance and cell area pulses predominantly exhibiting ∼10 mins period.

## INTRODUCTION

Semi-flexible biopolymers such as microtubules and actin filaments play important roles in cell signaling, cell division and cell motility within living organisms^1,2^. Physiologically, microtubules and actin filaments continuously hydrolyze adenosine triphosphate (ATP) or guanosine triphosphate (GTP) into adenosine diphosphate (ADP) or guanosine diphosphate (GDP) respectively^1,2^, while they undergo several stochastic transitions between phases of growth and shortening. They can be maintained in dynamic steady states under favorable kinetic conditions and in the presence of other regulatory proteins that modulate their dynamic characteristics including their assembly and disassembly^1,2,3,4,5^. They may form complex networks that spontaneously evolve into dynamic patterns, these exhibit stochastic fluctuations through changing numbers of nodes with dynamically varying biopolymer lengths due to growth and shrinkage and may undergo global or local network deformations^5,6,7^.

The formation of organized patterns observed across biological systems was first proposed by Alan Turing in his pioneering work using an activator-inhibitor model^8^. Gierer and Meinhardt^9^ developed similar models that have patterned solutions, in largely stationary and spatially homogeneous steady states. Small perturbations such as spatially heterogenous diffusion-driven instabilities induced pattern forming processes, which temporally evolved to stationary ‘Turing patterns’^8,9,10^. In the simplest Turing-type models, the inhibitor diffuses more rapidly than the weakly diffusive activator to limit the expansion of the activation. One can also arrive at this model by assuming that the activator is self-enhanced by consumption of a rapidly diffusing substrate. Both versions lead to formation of static periodic patterns like spots, stripes or labyrinths based on differential diffusion rates of interacting entities and the system parameters^11,12^. The presence of mechano-chemical feedback loops can drive such systems into far-from-equilibrium states leading to pattern dynamics^12,13,14^. Biological systems that exhibit patterning processes may involve several interacting entities with numerous unknown variables. The complexity of deciphering the key molecular players within such processes may vary spatiotemporally, particularly within in vivo systems. The key challenge then becomes extracting spatial-temporal evolution of dynamically patterned structures from experiments considering the bio-physical settings^15,16^. However, very few works exist towards quantifying motility and thereby the dynamics of molecular components within such networks to gain detailed understanding of the underlying principles of pattern formation.

Here, we focus on microridges, a special class of actin-based protrusions found on a variety of squamous epithelial tissues in several vertebrate species including fish-epidermis^17,18,19,20^. Morphologically, microridges typically grow into 2-100 µm lengths, forming labyrinthine patterns on cell surfaces and remain dynamic^21,22,23^, mimicking a class of reaction-diffusion ‘active Turing patterns’^22^. They undergo several stochastic transitions on minute timescales, presumably due to the underlying remodeling of actin network, resulting in their continuous re-organization^21,23,24,25,26,27^. These stochastic transitions occur over the course of embryonic development and later larval stages through multiple and simultaneous merging and splitting (or fusion and fission respectively) between microridges within a neighborhood^25,26,27^. However, how these events are dynamically regulated is not well understood. Recently, the molecular determinants within the microridges^22,23,25,28,29^ were uncovered and it has been shown that the pattern outcomes are considerably influenced by components of the polarity pathways^23^. However, due to lack of methods that can follow the precise motion of microridges, the dynamics of their formation, maintenance and the dynamics of proteins within them remains unknown.

Tracking and motion analysis are important tools for extracting quantitative information from live cells and organisms to gain insight into the dynamical and mechanical properties. Real time-lapse fluorescence microscopy images of various types of cellular motion or intracellular structures including regulatory proteins have drawn a broad range of tracking solutions that are largely custom-built, depending upon sizes and shapes of the imaged objects and the nature of motion of segmented objects^15^. A mathematical framework is constructed to solve the assignment problem to first link particles between consecutive frames and subsequently include gap closing depending upon the biological context^15,30^. In contrast to the tracking algorithms reported for dividing nuclei^31^, vesicle fusion/fission events^32^ or microtubule plus-end^33^; tracking of actin networks can pose severe challenges and requires specialized approaches. A skeleton image (after segmentation) of actin projections often contains several curves merging or splitting at a junction within an evolving network. How these merging or splitting segments are consistently followed temporally considering the topological complexities needs to be addressed appropriately during tracking. Few related works include multi-scale line detection and merging quasi-straight filaments^34^ or using ‘Stretching Open Active Contours’^35^ method for filament skeleton segmentation, followed by tracking skeletons using a k-partite matching approach^16,35^. Flow-based techniques^36^ compute average sub-region velocities^26^, but cannot provide positional information adequate for tracking. Recently, convolutional neural networks^26,37^ (CNNs) have considerably advanced detection and have been adapted for segmentation of biological data. However, finding an optimized solution for data association (linking trajectories) is the most challenging problem in tracking, which is largely solved using traditional computer vision methods^38,39,40,41^.

Here, we implemented a CNN-based framework for microridge pixel-detection and segmentation^26^, whose outputs were utilized for deriving the STIPS (*S*patio*T*emporal *I*nformation on *P*ixel *S*ubsets) algorithm. Our algorithm successfully solved the pixel associations by first building intermediate spatiotemporal pixel-mapping and then linking subsets of microridge pixel trajectories across consecutive time frames. From the STIPS outputs, we computed the fission and fusion event dynamics within a group of neighboring cells. Our analysis provided evidence for two distinct actomyosin based oscillations within neighboring cells; a common rhythmic activity governing both fusion and fission events directly contributing to maintenance of microridge patterns, apart from a coordinated apical cellular-level pulsatile activity.

## RESULTS

### Pre-trained CNN for pixel-wise microridge detection

A frame-by-frame cell-centroid-distance tracking algorithm^26^ was implemented to follow the spatial-temporal evolution of microridges within the same cell (Methods). The temporally ordered 2D-grayscale images of microridges within cell center regions of size 100×100 pixels were passed through a pre-trained microridge detection network resulting in microridge pixel-wise detection^26^ (Methods, Figs 1a-b). The starting point for the STIPS approach was the output unit activation of the ‘SoftMax’ layer whose values represented the maximum likelihood classification or detection probabilities, *ρ*^*ij*^ (values between [0,1]), and *ij* are the 2D-cartesian indices of each pixel within an image frame. We applied a threshold of t0.5 such that the detection probabilities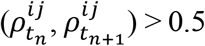, identified all the foreground pixels while the remaining values formed the background pixels over two consecutive 2D images frames, 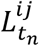 and 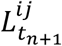 at t_n_ and t_n+1_ respectively (Fig 1c), similar approach proposed by Newby and colleagues^4^. The tracking challenge addressed in this work across consecutive time frames is illustrated in Fig 1d, showing green pixels at t_n_ that are shifted to new neighborhood locations (magenta) at t_n+1_, with common foreground (white) and background (black) pixels (Movie S1). Here, we present the STIPS algorithm that enabled consistently following the complex spatial-temporal evolution of microridge topology (Fig 1e).

**Fig 1.**
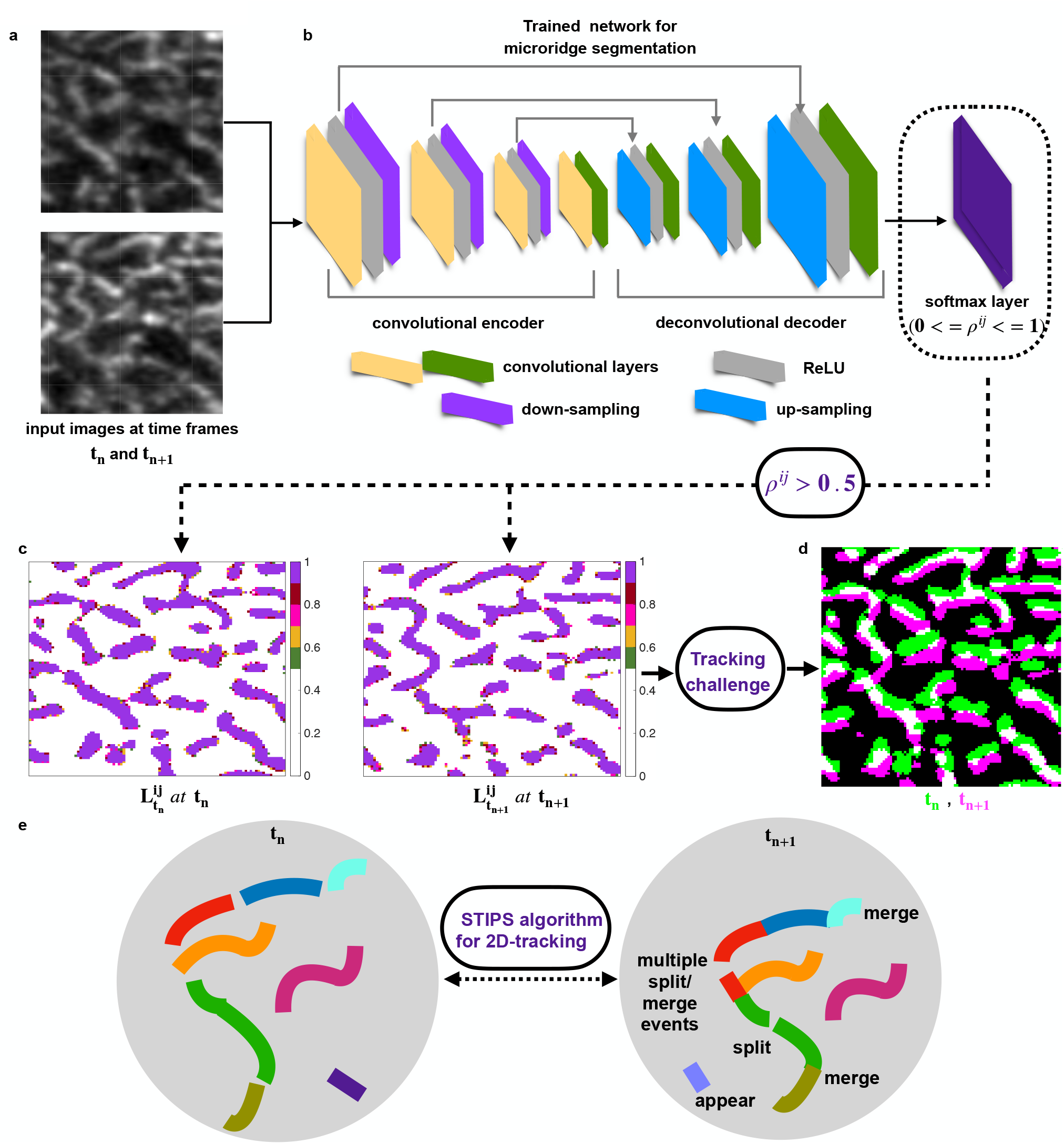
Pixel-wise detection of microridges and challenges in microridge tracking. **a**. Input 2D grayscale image sequences of microridges over consecutive time frames, t_n_ and t_n+1_. **b**. Pre-trained microridge segmentation network resulted in pixel-wise segmentation of all image sequences. ‘SoftMax’ layer stored the detection probabilities *ρ*^*ij*^, where *ij* are the indices for each pixel within a single time frame. **c**. By applying a threshold (*ρ*^*ij*^ *>0*.*5*) we identified the foreground pixels and all values below the threshold formed the image background (or 0) for each time frame, with their pixel coordinates 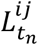 and 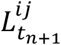 at t_n_ and t_n+1_, respectively. **d**. Tracking challenge across consecutive time frames with overlapping pixel regions in white, green pixels only existing at t_n_ and magenta ones only at t_n+1_ addressed in this work. **e**. Schematic representation of the challenge for the STIPS (*S*patio*T*emporal *I*nformation on *P*ixel *S*ubsets) algorithm for linking trajectories across consecutive time frames.

### Framework for pixel-wise changes across consecutive time frames

The growth or shrinkage events are purely stochastic and hence no pre-defined rules of interactions were set a priori across time frames for linking microridge pixel trajectories. The image frames, 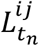 and 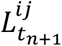, were converted into their respective binary images 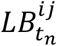 and 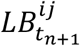, representing background and foreground pixels as 0 and 1 respectively (Figs 2a-b). We then performed a pixel-wise mathematical operation between 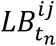 and 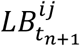 that uniquely identified the ‘state change’ of each pixel from frame t_n_ to t_n+1_, given by,

**Fig 2.**
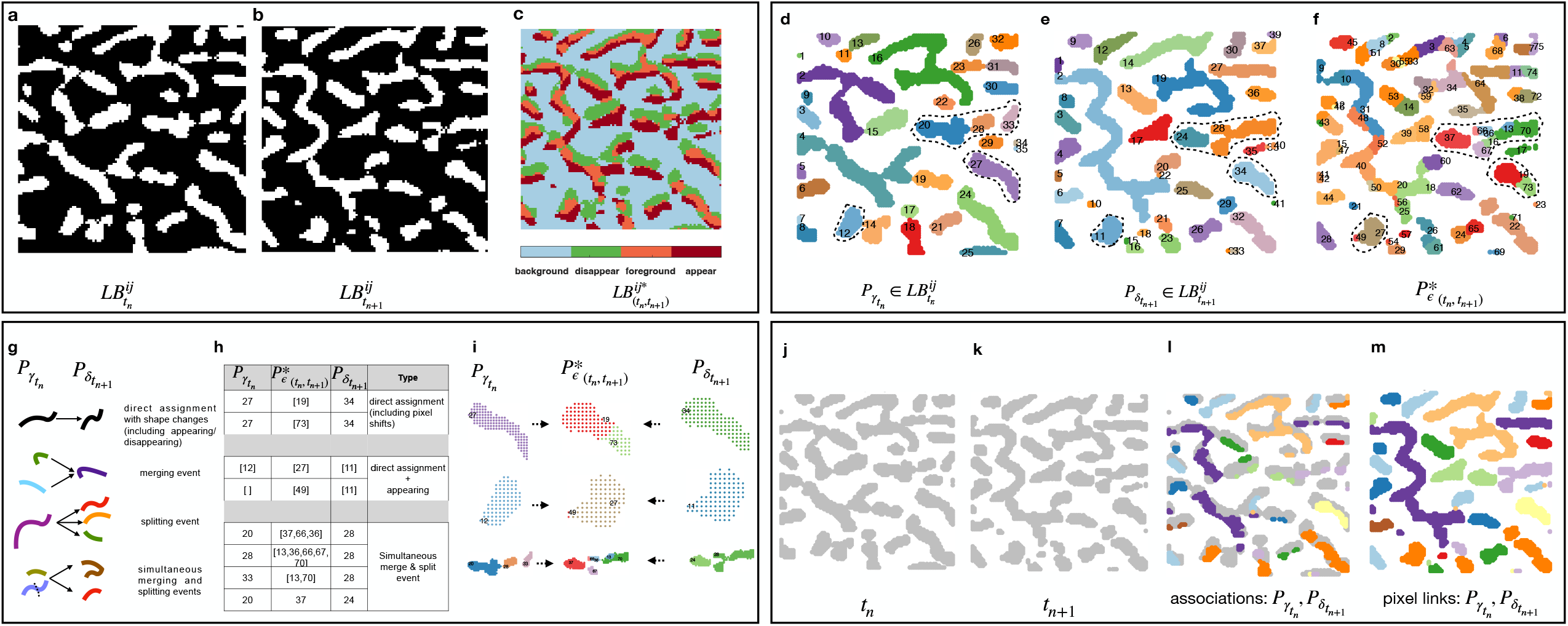
STIPS algorithm for solving microridge pixel-wise spatiotemporal associations - across consecutive image frames. **a-b**. Binary images 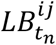 and 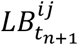 representing 0 for background pixels and 1 for foreground pixels respectively. **c**. Pixel-wise mathematical operation across consecutive image frames, 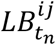 and 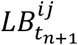 uniquely identified the four different ‘state changes’ of each pixel from frame t_n_ to t_n+1_, namely common foreground or background pixels across time frames and ‘appearing’ and ‘disappearing’ pixel transitions from t_n_ to t_n+1_. **d**. Connected sets of pixels for each microridge, 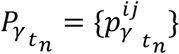 at frame t_n_ for image frame 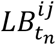 are shown in different random colors for each set, also displaying the set index *γ*. **e**. Similarly, the connected set of pixels of each microridge at 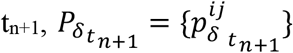 such that (*δ =1, 2*, …, *Δ*) are shown in different colors. **f**. Structure matrix 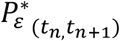 stored pixel associations derived from 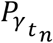 and 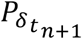 by sub-dividing pixels into mixture of cluster subsets (shown in distinct color), considering all possible merging and splitting events across consecutive time frames. All colors in **d. - e**. are arbitrarily chosen to show pixel subsets only and do not indicate any associations across time frames. **g**. Possible associations include pure merging or splitting events, multiple microridges simultaneously merging and splitting or direct one-to-one assignments, including appearances and disappearances of pixel coordinates between connected sets γ and δ. **h**. Transition table rows are examples from the data showing prospective microridge links across consecutive frames, with pixel subsets labeled γ at t_n_ linked to pixel subsets labeled δ at t_n+1_ using the intermediate array 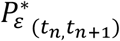. Three types of pixel re-arrangements across time frames are shown, also highlighted as dotted regions in **d. - f. i**. Using spectral clustering on the transition table, trajectory links for pixel subsets were resolved to map the associations between pixels from their coordinate lists across time frames. **j. - k**. Microridges in grayscale after segmentation at time frame t_n_ and t_n+1_ respectively. **l**. Microridges after segmentation at t_n_ (grayscale) superimposed with regions of identically colored pixels according to their final association with t_n+1_ (matching colored pixel regions or landmark pixels with microridge pixels at t_n+1_). **m**. Final results of the STIPS algorithm that builds pixel-wise spatiotemporal associations to establish microridge pixels subset trajectory links across time frames. Newly appearing pixels were appended whereas disappearing pixels at t_n_ are not colored. Matching pixel colors demonstrate associations across time frames in **l**. and **m**., demonstrating that the STIPS algorithm correctly links the microridges frame-to-frame.

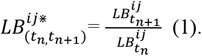

The matrix 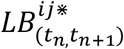 (Fig 2c) contained four distinct values, which labelled pixel state changes across consecutive frames in our Matlab implementation: (i) a pixel that remained background in both frames resulted in ‘0/0’, represented as ‘NaN’, (ii) a foreground pixel in t_n_ that disappeared at t_n+1_ resulted in ‘0/1’ or ‘0’, (iii) all the foreground pixels common across time frames remained ‘1’, (iv) pixels that were background at t_n_ but changed to foreground at t_n+1_ resulted in ‘1/0=Inf’ representing newly appeared pixels at t_n+1_. The four distinct ‘pixel-wise state changes’ facilitated building sub-trajectory links across consecutive frames.

### Connected set of pixels with their spatial coordinates form microridges

Using the method of connected components on the binarized image frames 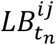 and 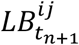 respectively, the connected sets of foreground pixels were gathered as single microridge along with their 2D spatial pixel coordinates. Each connected set of pixels were assigned unique identification numbers (ID’s) separately at each time frame. Specifically, for a total number *Γ* microridges at t_n,_ the connected pixel sets within a microridge (*γ*) were denoted by 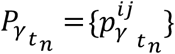 where *γ =1, 2*, …, *Γ* and *p*^*ij*^ represented the 2D spatial coordinates of each connected set that formed a microridge *γ* (Fig 2d). Similarly, let *Δ* be the total number of microridges at t_n+1_ and the connected pixels be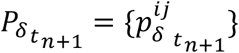, where *δ =1, 2*, …, *Δ* (Fig 2e). The number of connected components varied across frames, as did the number of 2D coordinates from one microridge to another within a time frame.

### Setup for solving spatiotemporal associations

In order to solve the pixel-wise spatiotemporal association problem, we initialized a structure matrix 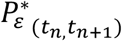 derived from pixel coordinates of 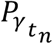 and 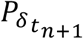 to trace changes in coordinate positions across consecutive frames. We assigned pixel coordinates 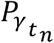 of 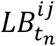 to 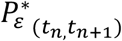 given by,

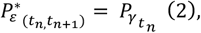

such that

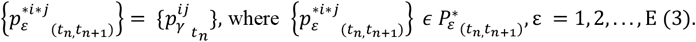

We identified the 2D positions assigned as 0 in the image matrix 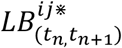 and removed such pixel coordinates from 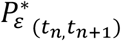, thus deleting foreground pixels at t_n_ that disappeared at t_n+1_ to flag either splitting events or shear movements, given by,

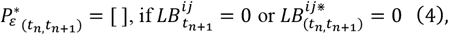

such that 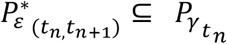 for microridge γ, containing common foreground pixels only across t_n_ and t_n+1_.

We obtained all spatial coordinates assigned as ‘Inf’ within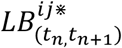, which accounted for newly appeared pixels at t_n+1_, where the key decision was which specific connected set *γ* at t_n_ to assign these to, or whether to create a new set at t_n+1_. Next, all such 2D coordinates 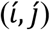 were identified and an exhaustive nearest neighbor search object model (Mdl.X) was implemented, where X is the query coordinate list of pixel positions 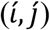 and the search space was each connected set 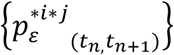 in 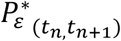.

Find 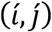 such that:

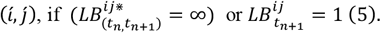

For each 2D pixel query coordinate, the model object (Mdl.X) performed an exhaustive search within 2-pixels radius (empirically determined maximum linking distance) to append query coordinates, 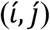 to the nearest existing foreground pixel 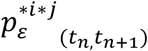 within 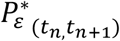. This approach merged newly appeared pixels at t_n+1_ into existing foreground pixel subsets such that 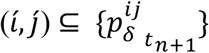.

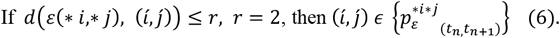

We then implemented a 2-step ‘density-based spatial clustering of applications with noise’ (DBSCAN) to find non-linearly separable clusters. For all the remaining unmatched query Points 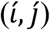, we first probed for the presence of any sub-clusters (minimum 1-point) within the query points, and all valid neighbors within 1-pixel radius (empirically determined) were clustered together and appended to 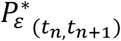 as separate coordinate lists. A second DBSCAN pass within 1-pixel search radius was implemented on coordinate list of 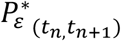 to check for presence of any sub-clusters within each coordinate set, to either form new disjoint clusters by splitting each set, or remain as connected coordinates otherwise. The result of this merge-split approach sub-divided pixels into several disjoint clusters represented by the array, 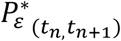 (Fig 2f). These steps considerably reduced the cost of forming pixel subset trajectories and avoided false positive links across time frames.

### Building transition table across consecutive time frames

The 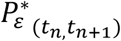 array (Fig 2f) captured all possible prospective associations between pixel subsets of γ at t_n_ with δ at t_n+1_ for solving the two-frame pixel-wise assignment linking problem (*Γ*≤ *Δ or Γ*> *Δ*). The possible associations included direct assignments, single or multiple merging/splitting events along with co-occurrences of appearances and disappearances of pixel subsets across consecutive frames. For each connected set γ at t_n_, we identified the associated connected sets of ε coordinates within 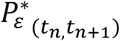 using set intersections (Figs 2d, 2f). Similarly, each ε coordinates was also associated with each δ-pixel coordinates at t_n+1_ (Figs 2e-f). The transition table was constructed between the γ-pixel subsets at t_n_ and δ at t_n+1_ using the common ε-connected sets (prospective microridges) that provided an association map for building sub-trajectory links (Figs 2g-h).

### Linking pixel subsets across consecutive time frames using the transition table

The transition table provided microridge links between γ at t_n_ and δ at t_n+1_, but did not indicate which exact pixels of γ at t_n_ were associated with the pixels of δ at t_n+1._ Hence, to re-assign the spatial coordinates across time frames, the trajectory links were resolved case-by-case to handle all possible scenarios to obtain correct pixel associations: direct one-to-one associations, appearances or disappearances and single or multiple merging/splitting events (Figs 2h-i). The transition table provided landmark pixel associations between pixel subsets of 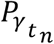 with 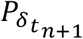 to construct trajectory links after transient re-arrangements within a local neighborhood region.

1. The simplest cases were direct one-to-one assignments (between γ and δ) with variations in pixel positions due to microridge movements and these were allocated to their nearest neighboring connected sets.
2. The empty arrays at t_n_ or t_n+1_ indicated disappearances and appearances of γ or δ respectively and hence such associations were assigned accordingly using a blank index by either terminating the γ-subset or starting a new δ-subset to be tracked from the next time frame.
3. There were either pure merging events (flagged by multiple pixels of γ-subsets associated with the same δ-subset) or pure splitting events (single γ-subset associated with multiple δ-subsets) or simultaneous merging and splitting events (multiple γ-subsets associated with several δ-subsets), including concurrent pixel appearances and disappearances in all cases. As a rule, merging events were resolved by concatenating pixels of γ-subset lists at t_n_ transiting into δ-subsets at t_n+1_, whereas splitting events were solved by segregating γ-subsets into the associated number of δ-sub-clusters at t_n+1_. We concatenated pixel coordinates labeled γ or multiple γ’s [γ_1_, γ_2, …_ γ_q_], merged together as a single connected set at t_n_. This was followed by resolving the splitting events using a similarity graph model, whose inputs were the concatenated γ-labeled pixel coordinates that were split into *p*-clusters for [δ _1_, δ _2_ … δ_p_], where *p* represented the associated δ’s at t_n+1_. A 3-pixel search radius was set for assigning the δ-sub-clusters to their nearest γ-sub-clusters using spectral clustering based on the minimum Euclidean distances between the mean centroids of the sub-pixel clusters. For uneven distribution of pixel coordinates along x versus y-directions, the pixels were normalized with their standard deviations (> 3.5 pixels) prior to spectral clustering (Figs 2h-i).

The STIPS algorithm takes two consecutive frames as inputs (Figs 2j-k) and creates pixel subset associations between them (Fig 2l) to establish pixel-wise spatial-temporal links frame-to-frame (Fig 2m), demonstrated across 59-time frames (32.8 mins) (Movie S2).

### STIPS algorithm yields temporal evolution of microridges while accounting for fusion and fission events

The most striking features of microridge dynamics are the fusion and fission events involving merging and splitting of two or more microridges^22,25,26,27^. The STIPS algorithm enabled microridge tracking within entire periderm cells of zebrafish embryos at 2-dpf (days-post-fertilization), when microridges are highly dynamic (Movies S3-S4). From these tracks, we computed the temporal sequence of fusion and fission events. When the same microridge label at t_n_ was associated with multiple microridges at t_n+1_, this was considered fission. Multiple microridge labels at t_n_ that corresponded to a single microridge at t_n+1_, were identified as fusion events. We determined the total number of microridges (black line, Fig 3a) at each time point and found that the total numbers only showed small variations with respect to the temporal mean of number of microridges (green dotted line, Fig 3a). However, the typical number of fissions and fusions (in blue and red stem lines respectively, Fig 3a) fluctuated around the mean number of microridges. Most interestingly, the temporal evolution in both event numbers exhibited cyclic changes leading us to probe the presence of any prominent oscillations in their event dynamics.

**Fig 3.**
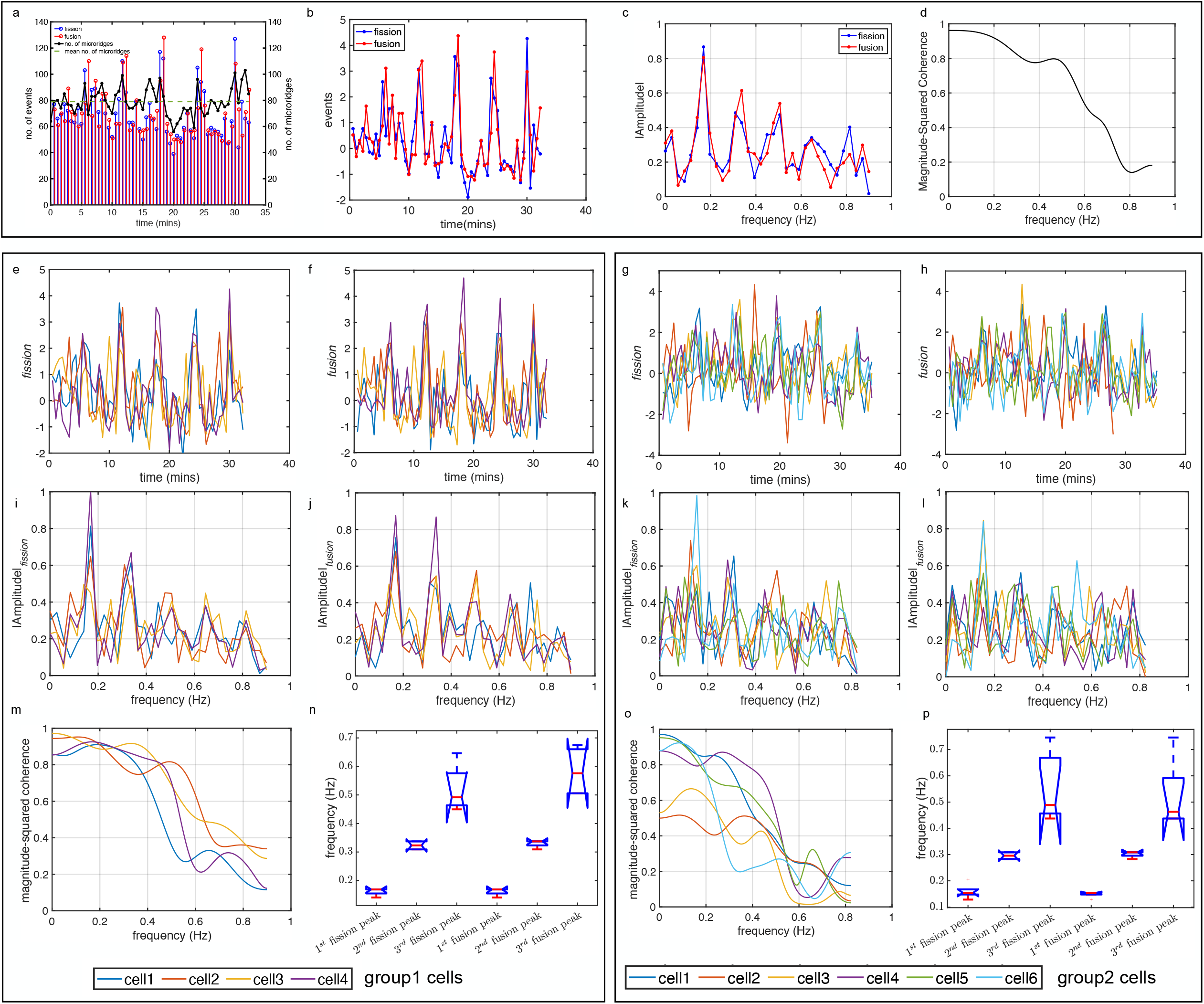
Similar prominent frequencies in fusion and fission event time series within a cell and coordinated pulses across a group of neighboring cells. **a**.Number of fission and fusion events (in blue and red stem lines respectively) across consecutive time frames. Total number of microridges (black line) at each time point within a yolk cell (Movie S3) fluctuated close to the mean number of microridges (green dotted line). **b**. Fusion and fission event numbers normalized with their median values and median adjusted deviations indicated concurrent peaks. **c**. Fast Fourier transform (FFT) of the time-domain signal provided the single-sided amplitude spectrum at a sampling frequency F_s_= 1.79 Hz, that exhibited three distinct frequency peaks in the amplitude signal for both fission and fusion events respectively. **d**. Magnitude-squared coherence between the fission and fusion event signals (median normalized) indicated strong coherence ( > 0.8) for small frequencies (or longer periods) upto 0.35 Hz (∼2.85 mins period) and less coherence for higher frequencies (or shorter periods). **e. - h**. Time domain fusion and fission events signal (median normalized) for a group of 4 neighbouring cells (group1 cells) and 6 neighbouring cells (group2 cells) respectively of the yolk region exhibiting noisy oscillations. **i. - l**. Single-sided spectrum of fusion and fission events signal computed at a sampling frequency, F_s_= 1.79 Hz for group1 cells and F_s_ = 1.65 Hz for group2 cells respectively. Three distinct prominent frequencies within similar range for both the events observed for all the cells within both the groups of neighbouring cells. For group1 cells, the three significant amplitude peaks occurred between 0.14 and 0.16 Hz, 0.30 to 0.33 Hz and 0.44 and 0.67 Hz for both fission and fusion events. For group2 cells prominent amplitude peaks occurred between 0.12 to 0.15 Hz, 0.28 to 0.30 Hz and 0.43 Hz to 0.74 Hz respectively for both the events. **m**., **o**. Strong coherence between fusion and fission event signals at lower frequency values that steeply decreased for all the cells for higher frequency range for both groups, with relatively greater variance observed in group2 cells. **n**., **p**. Box plot showing three prominent frequencies and their variance in the ensemble of all observed cells; shorter frequency cycles (longer time periods) exhibited a smaller variance (1^st^ and 2^nd^ peaks) in comparison with higher frequency values or shorter time periods (3^rd^ peak) within both the groups.

### Fission and fusion events shared common dominant frequencies indicating significance of actomyosin based oscillations

In order to determine the dominant frequencies in the fusion and fission events time series, within a cell, these were normalized with their temporal median values (Methods, Eq. 1-2) that evidently showed synchronized recurring peaks for both types of events emerging among noisy fluctuations (Fig 3b). To assess the dominant frequencies in these time series, we computed the frequency domain amplitude spectrum of the two signals. We computed the fast Fourier transform (FFT) of the time-domain signal for the fusion and fission events separately for a selected cell and amplitude peaks were identified for both the events (Fig 3c). The amplitudes peaked at similar prominent frequencies for both fission and fusion events. The lowest prominent frequency was at 0.16 Hz (5.9 mins period) for both events, while the second and third peaks occurred at 0.33 Hz and 0.5 Hz (2.9 and 2 mins period) respectively for the two events (Fig 3c). To further elucidate that frequencies of oscillations are common to both event time series signals, we estimated the frequency-dependent cross-spectrum (Methods, Eq. 3) that indicated strong coherence (>0.8) over small frequencies (or longer periods) upto 0.35 Hz (∼2.85 mins period), which steeply decreased for higher frequencies largely owed to signal fluctuations (Fig 3d).

We then sought to examine microridge patterns within two different groups of neighbouring cells from the yolk region (Figs 3e-p). The time domain fusion and fission event counts (normalised) for group1 (4 cells) and group2 (6 cells) exhibited similarly peaked, however noisy oscillations (Figs 3e-h). Strikingly, the amplitude spectrum exhibited at least three distinct prominent peaks at very close frequency between fission and fusion events for all the cells within each group (Figs 3i-l), lending support to the hypothesis that a common biophysical mechanism is responsible for them. There were indications of strong coherence between fusion and fission events for lower frequency values (upto ∼0.35 Hz for group1), that steeply dropped with increased frequency, comparably also observed in group2 (Figs 3m, 3o). The 1^st^ and 2^nd^ peaks exhibited a smaller variance, while with higher frequency values, greater jitters in the spectrums were visible for both the events within each group that could be attributed to biological noise as well as due to cycle to cycle fluctuations in (instantaneous) amplitude and (instantaneous) period (Figs 3n, 3p). Similarly, peaks in fusion/fission events for group3 (4 cells) within the head region (Fig S1), was in qualitative agreement with groups1-2. Our results suggested a common oscillatory mechanism coordinated across neighboring cells governs both the events.

We also observed pulsatile activity in cellular area measured from the raw data of periderm cell image timeseries within the three groups and examined whether these were associated with prominent fusion or fission amplitude peaks. The fission and fusion signals occasionally exhibited peaks or troughs coherent with the normalized area temporal signals (Methods, Eq. 4, Figs 4a-c, S2a). The area frequency spectrum indicated a single strong amplitude peak at ∼0.1 Hz (10 mins period) followed by low amplitude fluctuations across the remaining frequency spectrum for all cells within the three groups (Figs 4d-f, S2b). Consequently, coherence was also found to be low between area versus fusion and area versus fission (Figs 4g-h, S2c), indicating that the underlying frequencies responsible for area fluctuations rarely coincided with fusion/fission events (Figs 4g-h). Next, we examined the net cell area changes due to their pulsatile activity (Methods, Eq. 5). We observed varied behaviors; small-scale fluctuations, over-all expansion or net cellular contractions (Fig 5a), however the frequencies underlying fission and fusion event cycles remained independent of area frequency, in contrast to the frequency coherence between fission and fusion within all cells across the three groups (Fig 5b).

**Fig 4.**
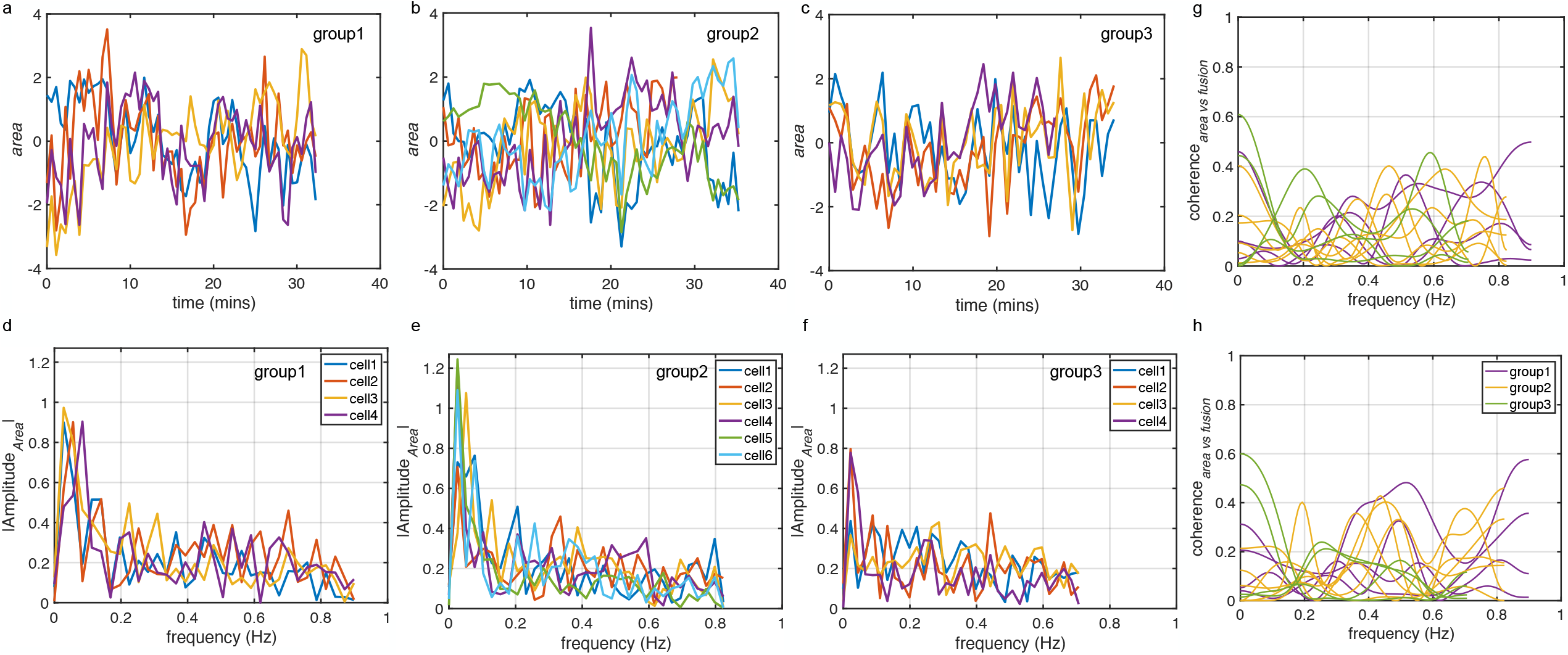
Dominant frequency in area pulses did not coincide with fission/fusion frequencies within the three groups of cells. **a.-c**. Time domain area (median normalized) of neighbouring cells within the three group. **d.-f**. Single-sided amplitude spectrum in the Fourier domain indicated a distinct amplitude peak at ∼0.1 Hz for all the cells within the 3 groups, followed by several degrees of irregular frequencies. **g.-h**. Magnitude square coherence of area versus fusion and area versus fission event computed for within a cell exhibited low values (consistently observed across all three groups) respectively.

**Fig 5.**
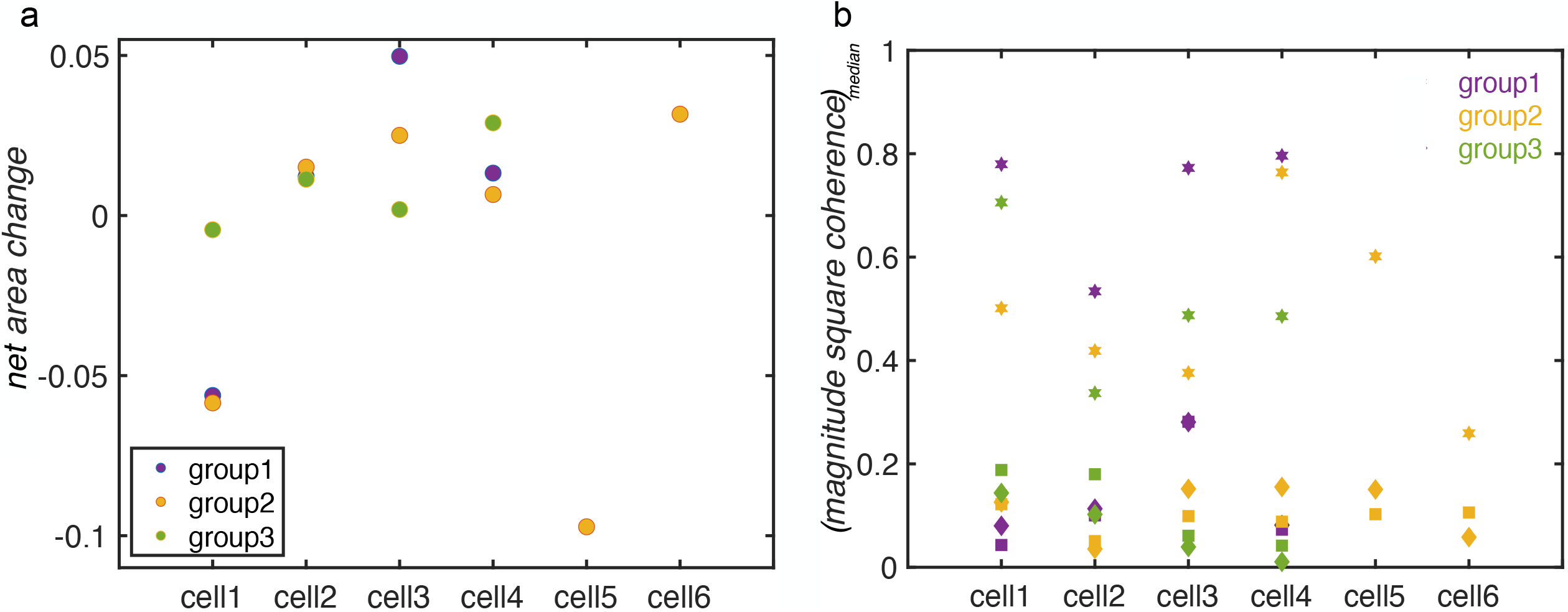
Fusion and fission events are independent of net area changes. **a**. Changes in net area across all cells within the three groups indicated varied behavior; nearly constant, net increase or net decrease. **b**. Magnitude square coherence (represented by the median values of the curve) between fission and fusion (star) is relatively high for all cells across different groups, whereas the coherence between fission and area (diamond) and coherence between fusion and area (square) respectively remained below 0.3 for all cells within the three groups.

In summary, our STIPS approach enabled tracking the complex dynamics of microridges and evaluating fission/fusion event dynamics. Our analysis elucidated cell-group rhythmic dynamics at two different levels; a common pulsatile activity with three prominent peaks between 2 and 5.9 mins periods governing both fusion and fission events contributing to pattern dynamics and an evident apical cellular area pulses with a dominant ∼10 mins period.

## DISCUSSION

Several complex self-organized patterns have been explained using Turing-type mathematical models with few biologically tractable entities (constants within the model are molecularly significant) to provide insight into mechanisms of patterning processes. However, often it becomes challenging to delineate biochemical, genetic and mechanical regulators acting in a spatiotemporal manner over longer developmental timescales. The microridges are one such example that consist of actin filaments, myosin motors, various cross-linkers and actin-binding proteins ^20,22,23,24,25,28,29^, which are dynamically formed and maintained from 1-dpf onwards upto 4-dpf and possibly regulate recurring fusion and fission events^25,26^ as a result of integrated responses of multiple proteins. Furthermore, the relative dominance of control mechanisms might dynamically vary over the course of embryonic development.

For instance, relative dominance of key molecular components might dynamically vary over the course of embryonic development. The interplay of key regulators of microridge formation during the early stages (∼1-dpf) might be different from key molecular components involved during microridge maintenance phase at later larvae stages (beyond 1-dpf upto 4-dpf). The dynamic properties of the interacting components within the microridges and their collective behavior are important for understanding their role in maintaining epithelial function.

Here, we have presented a novel and tractable approach to assess microridge dynamics by automatically tracking complex microridges flows. The STIPS algorithm enabled investigation of fusion and fission event dynamics, which revealed coherent beat frequencies within a cell. Surprisingly, we uncovered the presence of coherent beat frequencies within a group of neighbouring cells, indicating that the fusion-fission dynamics are regulated through rhythmic activation of cortical oscillations that are coupled (nearest-neighbour) in a frequency-dependent manner^42^. There were also indications of oscillations with different degrees of regularity facilitating small-scale spatial-temporal fluctuations contributing to maintenance of microridge patterns. Conceivably, diverse array of pulsatile activities within the epithelial cortex influence microridge patterning. A plausible explanation for frequency-dependent coupling of oscillations in fusion and fission events could be the pulsing duration of F-actin itself, as a consequence of NMII activity. It was previously reported that the apical domain of periderm cells exhibited pulsatile apical contractions of the actomyosin cortex via NMII pulses, necessary for the initial formation of microridges^24^ and that the local NMII activity was directly associated with fission and fusion of microridges^25^. Increasing the levels of pMLC by over expressing constitutively active Myosin Light Chain Kinase (MLCK) resulted in increased microridge lengths and more complex pattern^23^, whereas inhibiting NMII activity by using Blebbistatin resulted in the breakdown of microridges^23,28^. On the other hand, inhibiting ROCK decreased microridge lengths and significantly reduced NMII pulses, indicating that the activity of RhoA drives NMII contractions of the apical actomyosin cortex^24^. In several systems, the spatiotemporal pattern of cortical contractility in a pulsatile manner can be achieved through RhoA GTPase signalling pathway, which activates NMII through the regulation of Rho-associated kinase (ROCK) and MyoII light-chain phosphatase^23,24,25,43^. Here, we have shown that the rhythmic dynamics in fission and fusion cycles is important for maintenance of microridge patterns. Moreover, pulsed signalling decoded in a frequency-dependent manner enabled coupling among neighbouring cells to maintain these oscillations. Indeed, as a biological timer, pulsed signalling is better tuneable and adaptive than continuous signalling, particularly when temporal order is employed for dynamically maintaining spatial patterns over prolonged durations^42,44,45^.

We also evaluated pulsatile activity in cellular area that revealed a strong amplitude peak with a ∼10 mins period. Besides, cell area changes due to pulsatile activity was largely insignificant for fission or fusion events during microridge maintenance phase. Previously, it was reported that apical constriction is the main driving force for formation and elongation of microridges indicated by continual decrease in apical area and simultaneous fusion events, while increase in area concomitantly reduced microridge lengths (fission events) around 1-dpf^24^. Our results indicated that during maintenance phase, a common pulsatile mechanism drives both fusion-fission dynamics to progressively build and maintain a self-organizing patterning system, largely independent of cellular-level pulsatile activity.

Our approach is adaptable to similar dynamical systems undergoing temporal stochastic transitions in complex environments and importantly would enable motility measures of multiple proteins over extensive temporal/spatial scales. In future, methodical improvements on imaging technologies^46^ would substantially enhance spatial-temporal continuity at nanometer resolution to aid in tracking. Concurrently, deep-learning-assisted-tracking^47^ hold promise to revolutionize our understanding of complex dynamic patterning in the near future.

## METHODS

### Zebrafish strains

The experimental procedures were performed on zebrafish lines *Tg(actb1:GFP-utrCH)* gifted by Behrndt and colleagues^48^, which we also previously reported^26^. The zebrafish were maintained at standard laboratory conditions. The institutional animal ethics committee approved the experimental protocols used in this study.

### Mounting zebrafish embryos for live image acquisition

The 2-2.5dpf stages zebrafish embryos maintained in E3 buffer with 0.02% ethyl-m-aminobenzoate methanesulphonate (Triacane) were mounted on an agarose-free flat mounting setup, custom fabricated at the TIFR workshop, was used for live imaging the lateral side of the embryo. A 40x dipping lens upright Zeiss confocal 880 microscope recorded z-stack images from apical upto basal epidermis at time intervals of 0.55 or 0.60 mins of the cells from yolk regions^26^. The images were acquired with lateral pixel size 0.1977 µm and axial pixel size 0.5-0.6 µm, respectively at a preset room temperature with fluctuations between 25°C - 27°C.

### A custom-built image processing pipeline produces the training dataset for CNN based microridge segmentation

A two-step segmentation and tracking approach was implemented, as described previously^26^. Firstly, we built a custom method for cell membrane segmentation followed by cell tracking. Single cells were isolated with their microridges using cell boundary information. Cells were tracked by using the nearest neighbor cell centroid distances for frame-by-frame cell association and assignment. Single cell isolation and subsequent tracking allowed to follow the same cell over all time frames and thereby also follow its intrinsic microridge dynamics. For pixel-wise identification of microridge patterns, a second custom-built segmentation algorithm was designed with a series of image processing techniques to produce a segmentation mask of the microridges where each pixel was assigned a label for microridges and another label for background pixels^26^. All the custom-built codes are available at https://github.com/Bhavna-R/DeepLearningQuantAnalysActinMicroridges.

The pairs of extracted raw microridge cell images and their corresponding annotated images formed the dataset for training a convolutional neural network (CNN) for microridge segmentation. We implemented a semantic segmentation approach, which converts the image segmentation problem into a binary pixel-level image classification problem. The U-net network hyperparameters were optimised to achieve high pixel-wise segmentation. An image size to 256^2^ (pixels), that scales to the receptive field size with encoder depth of 6 and the learning rate as 10^-4^, mini-batch size equal to 6 and maximum epoch of 800 yielded ∼95% accuracy measure^26^.

### Microridge pre-trained CNN network for microridge pixel-wise detection within a cell time series

Cell tracking^26^ subsequently allowed following the microridge dynamics within the same cell. The grayscale temporally ordered cell images patterned with microridges were passed through the already trained microridge segmentation network. The output of the ‘softmax layer’ or the normalized exponential function which is just prior to the classification layer of the trained network formed the input to the STIPS algorithm. The temporally extracted cell images were part of either the training dataset or the test dataset that was used for obtaining the trained microridge segmentation network. Some grayscale cell images were not part of the dataset. The dataset setup for obtaining the segmentation network does not affect the construction of the STIPS algorithm.

### Determination of fusion and fission events across time frames

Considering two consecutive time frames, when the same microridge label at t_n_ was found to be associated with microridges at t_n+1_ at multiple times (>1), this was considered as fission event. Similarly, multiple microridge labels at t_n_ that corresponded to a single microridge at t_n+1_, indicated a fusion event. Each of the events were summed up across consecutive time frames that provided the total number of events across t_n_ and t_n+1_. The number of microridges was summed up to provide the total number of microridges at any given time frame (t_n_). To estimate the mean number of microridges, we computed the time average over the number of microridges for a given cell.

### Fission and fusion events timeseries analysis

In order to compare the events time series within any given cell or a group of neighboring cells, the fusion and fission events time series were normalized with their median values and adjusted with their median absolute deviation (mad). Let fission event and fusion event time series be denoted by *F*_*i*_*(t)* and *F*_*u*_*(t)* respectively, the normalized series are

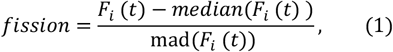

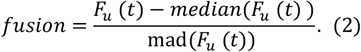

In order to compute the single-sided Fourier spectrum (using only positive frequencies) of the fission and fusion signals, the fast Fourier transform (FFT) was computed with sampling frequency F_s_= 1.79 Hz, 1.65 Hz, 1.41 Hz determined from the sampled imaging frame rate of 0.55 mins, 0.60 mins, 0.70 mins for group1, group 2 and group3 cells respectively.

The absolute signals of the event time series were normalized with their respective lengths and the DC component of the signal that corresponds to zero frequency was removed. The frequency bin was determined using Fs/N, where N is the length of the time series signal. The peak in the amplitude signals were determined automatically using the ‘*findpeaks*’ function in Matlab with a minimum peak prominence value set to 0.325 and the minimum peak distance set to the sampling frequency. Considering the jitters within the frequency spectrum, the three prominent peaks were manually assigned to the 1^st^, 2^nd^ or 3^rd^ peak category. This was due to low amplitude peaks leading to missing frequency peaks for some cells. To ensure that the peak signal observed in fission events is simultaneously observed in the fusion event or vice-versa, we computed the magnitude squared coherence (values between 0 and 1) that estimates how the two signals corresponded to each other at each frequency. The magnitude squared coherence between the fission and fusion signals was computed as a function of their power spectral densities, given by *P*_*fi*_*(f)* and *P*_*fu*_*(f)* respectively and their cross power spectral density *P*_*fi*,*fu*_*(f)*,

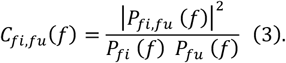

### Cell area temporal signal analysis

We used similar technique (as for fusion and fission events) to normalize cell area timeseries. The temporal area signal was directly obtained from raw cell images (prior to passing through the CNN) and the areas were normalized with their median values and adjusted with their median absolute deviation (mad):

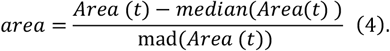

The sampling frequency was directly computed from each time series and the single-sided Fourier spectra (using only positive frequencies) were computed. The absolute signals of the event time series were normalized in the same manner as that of fusion and fission signals. The peaks in the amplitude signals were visually observed and verified for all the cells within the three groups. The magnitude-squared coherence was computed for each cell between its fission versus area and fusion versus area in the frequency domain within the different groups. The net cell area change for each cell was computed according to,

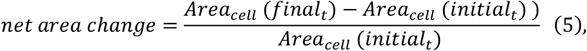

where, *final*_*t*_ and *initial*_*t*_ were the areas (µm^2^) at the final timepoint and initial timepoint respectively. The median value of the magnitude-squared coherence for each cell between its fission versus area, fusion versus area and fission versus fusion was chosen for an overall assessment for all cells across the groups.

## ACKNOWLEDGEMENTS

We acknowledge the department of atomic energy (DAE), TIFR for the funding (RTI4003;12P-121). We thank Sebastian Wüster for valuable comments and critical proofreading of the manuscript.

## AUTHORS CONTRIBUTIONS

RB- Conceptualization, methodology, investigation, data curation, formal analyses, visualization, writing manuscript draft and editing; MS- laboratory support for experiments, draft editing, revision, draft finalization.

## SUPPLEMENTARY FIGURE LEGENDS

**Fig S1. Oscillations in fusion and fission event numbers within a group of neighboring cells in the head region**

**a.-b**. Time domain fusion and fission event numbers (median normalized) within a group of 4 neighbouring cells (group3 cells) respectively of the head region. **c.-d**. Single-sided spectrum computed at a sampling frequency, F_s_= 1.41 Hz for the group3 showed three distinct amplitude peaks in the Fourier domains for all the cells within the neighbourhood for both the events. Significant amplitude peaks occurred between 0.04 and 0.08 Hz, 0.28 and 0.41 Hz and 0.30 and 0.66Hz for both events. **e**. Strong coherence between fusion and fission event signals observed at lower frequency values that decreased at higher frequencies. **f**. Box plot indicated low variances for the 1^st^ and 2^nd^ peaks in comparison with 3^rd^ peak that occurred at higher frequency values or shorter time periods.

**Fig S2. Cell area versus fusion and fission events variations**

**a**. Normalized cell area and fusion and fission event time series respectively for a single cell. Temporal peaks in fission/fusion events occasionally coincided with temporal area peaks/troughs. **b**. Single-sided spectrum indicated a single significant amplitude area peak at 0.1 Hz (∼10 mins period) followed by several low amplitude peaks. Fission and fusion events indicated three distinct amplitude peaks. **c**. Magnitude squared coherence between area versus fusion and area versus fission events remained low across all frequencies.

## SUPPLEMENTARY MOVIE LEGENDS

**Movie S1. Microridge tracking motivates specialized approach to follow the temporal evolution of their topology consistently**

Tracking challenge across consecutive time frames shows overlapping pixels that remain in white, green pixels only existing at t_n_ while disappearing at t_n+1_ and newly appearing magenta pixels at t_n+1_ addressed in this work. Microridge lengths and size are temporally altered and they continuously switch their connectivity with their neighbours via recurring fusion and fission events.

**Movie S2. Results of pixel-wise spatiotemporal associations across consecutive time frames**

Left panel indicates microridges in grayscale after segmentation at time frame t_n_ with matching colored pixels superimposed according to their mapping to microridge pixel subsets at t_n+1_. Right panel indicates microridge pixels subset trajectory links at t_n+1_ using the STIPS algorithm. Temporally linked microridges are displayed in the same matching colors in the left and right panels. Newly appearing pixels were also appended to the list in the right panel, whereas the disappeared pixels at t_n_ were removed from the list and are not displayed in colors in either panel. Two consecutive time frames from the movie are displayed in Figs 2l-m.

**Movie S3. Microridge tracking within a yolk cell (group1) using the STIPS algorithm** Representative movie from group1 cells, also shown in Fig 3a-d. Left panel indicates microridges in grayscale after segmentation at time frame t_n_ with matching colored pixels superimposed according to their mapping to microridge pixel subsets at t_n+1_. Right panel indicates microridge pixels subset trajectory links at t_n+1_ using the STIPS algorithm for the entire cell that was shown in Movie S2.

**Movie S4. Microridge tracking within a yolk cell (group2) using the STIPS algorithm** Representative movie from group2 cells. Left panel indicates microridges in grayscale after segmentation at time frame t_n_ with matching colored pixels superimposed according to their mapping to microridge pixel subsets at t_n+1_. Right panel indicates microridge pixels subset trajectory links at t_n+1_ using the STIPS algorithm for the entire cell.

