## Supplementary figures and images for "STIPS algorithm enables tracking labyrinthine patterns and reveals distinct rhythmic dynamics of actin microridges"

### Supplementary Figures

Fig S1. STIPS, Bhavna R, Sonawane M

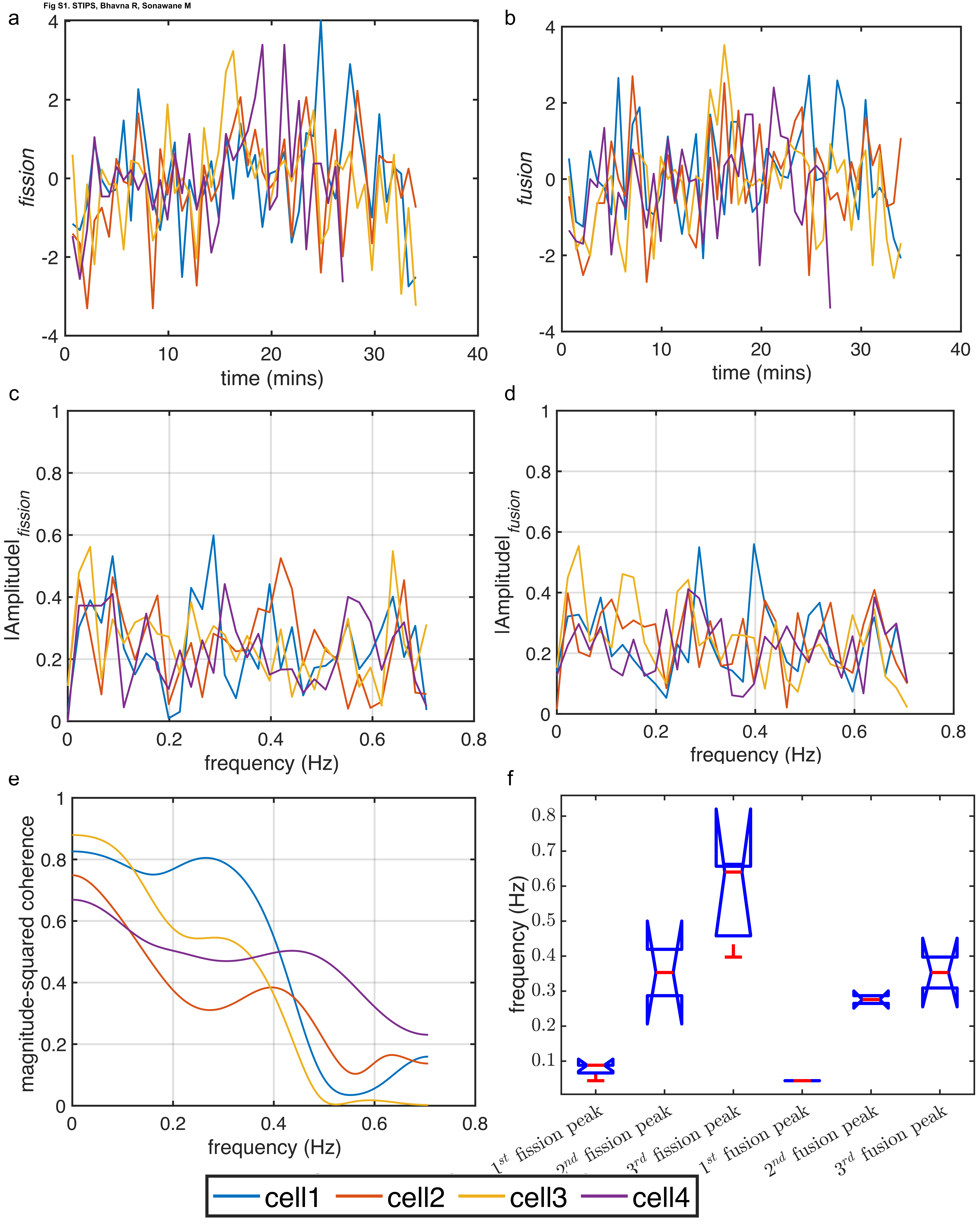

Fig S2. STIPS, Bhavna R, Sonawane M

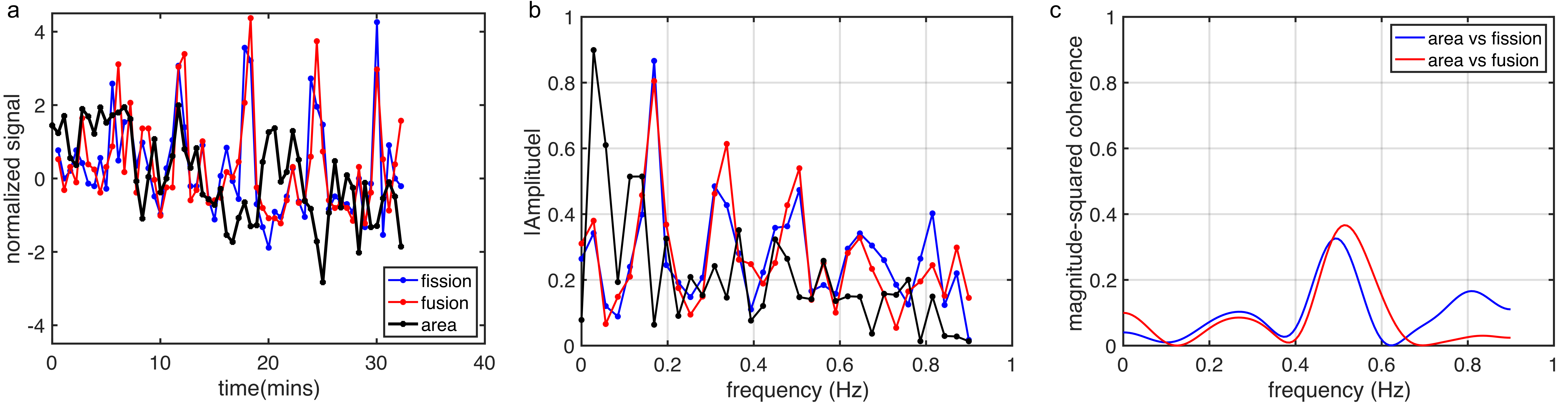
